# Novel, User-friendly Experimental and Analysis Strategies for Fast Voltammetry: 1. “The Analysis Kid” for FSCV

**DOI:** 10.1101/2021.04.08.439012

**Authors:** Sergio Mena, Solene Dietsch, Shane N. Berger, Colby E. Witt, Parastoo Hashemi

## Abstract

Fast-scan cyclic voltammetry at carbon fiber microelectrodes measures low concentrations of analytes in biological systems. There are ongoing efforts to simplify FSCV analysis and several custom platforms are available for filtering and multi-modal analysis of FSCV signals but there is no single, easily accessible platform that has capacity for all these features. Here we present *The Analysis Kid*: a free, open-source cloud application that does not require a specialized runtime environment and is easily accessible via common browsers. We show that a user-friendly interface can analyze multi-platform file formats to provide multimodal visualization of FSCV color plots with digital background subtraction. We highlight key features that allow interactive calibration and parametric analysis *via* peak finding algorithms to automatically detect the maximum amplitude, area under the curve and clearance rate of the signal. Finally, *The Analysis Kid* enables semi-automatic fitting of data with Michaelis Menten kinetics with single or dual reuptake models. *The Analysis Kid* can be freely accessed at https://analysis-kid.herokuapp.com/. The web application code is found, under an MIT license, at https://github.com/sermeor/The-Analysis-Kid.

---

Fast-scan cyclic voltammetry (FSCV) at carbon fiber microelectrodes is an electrochemical that measures low concentrations of analytes, with a good signal-to-noise ratio (SNR) and time resolution. Because of the micron dimensions of the electrodes, the technique has been popular for biological analysis, especially in intricate tissues such as the brain.^1^ FSCV has been widely applied to measurements of dopamine in *ex vivo* and *in vivo* models. Here, a triangular waveform is applied to the electrode at high frequency (10 Hz) and scan rate (~400 Vs^−1^).^2, 3^ The positive and negative limits of the waveform are chosen to optimize the signal per experimental preparation; for dopamine the limits are generally −0.4 V resting/negative potential limit (to facilitate preconcentration of the dopamine cation) and 1.3 V positive limit (to overoxidize the carbon surface thereby increasing sensitivity).^4^ Recently, detection waveforms have been optimized to measure serotonin, hydrogen peroxide, adenosine, copper and guanine, *in vivo* and *ex vivo*.^5–9^

FSCV is experimentally challenging, thus strong efforts have been in progress to simplify it. Several custom software platforms have been developed for the filtering and analysis of FSCV signals.^10–12^ These packages require memory-heavy runtime environments (such as LabVIEW’s) and are often not compatible with each other and/or between operating systems. This is particularly problematic because maintenance/updates and new analysis algorithms do not transfer between packages. Additionally, each package has a niche analysis feature (*e.g.,* Demon Voltammetry can model decay curves with Michaelis-Menten (M-M) kinetic models). There is no single, easily accessible platform that has the capacity for all these analyses.

Here we present *The Analysis Kid*: a free, open-source web application that does not require specialized runtime environments to combine the most useful aspects of FSCV analysis. The program is easily accessible *via* common browsers where post-processing of experimental signals with custom algorithms have minimal software dependency and maintenance and updates are automatic. A user-friendly interface can analyze multi-platform file formats (.csv, .txt, .xls and .xlsx) to provide multimodal visualization of FSCV acquisition with digital background subtraction.

In this paper, we first describe the background subtraction and filtering methods incorporated in our application (including 2D convolution smoothing and 2D fast Fourier transform (FFT) filtering). Second, we highlight key features that allow interactive calibration and parametric analysis *via* peak finding algorithms to automatically detect the maximum amplitude, area under the curve (AUC) and clearance rate (*t*_1/2_) of the signal. Finally, *The Analysis Kid* enables semi-automatic fitting of data with Michaelis Menten kinetics with single or dual reuptake models.

*The Analysis Kid* can be freely accessed at https://analysis-kid.herokuapp.com/. The web application code is found, under an MIT license, at https://github.com/sermeor/The-Analysis-Kid.

## EXPERIMENTAL SECTION

### Software

*The Analysis Kid* is designed to work as a lightweight front-end web application with minimal connection to the server. The graphical user interface was written in HTML, CSS, and JavaScript. The imported FSCV data and methods were organized using object-oriented programming. The application is hosted in a PHP runtime Linux server on a cloud platform. The server allows 256 users to simultaneously connect. Data processing algorithms were largely custom designed on JavaScript, except for the 2D FFT algorithm, compiled from C++ to JavaScript. The website was designed in a hierarchical structure. The applications can be accessed from a homepage, which contains documentation and tutorials on how to use them. This way, several applications can be opened from the homepage at the same time without any interference between them. The files are fully processed in the local desktop browser and are not loaded into the server. As a result, the files are safe in the user’s personal computer and the computational speed of the algorithms depends on the local machine.

FSCV data is graphed using the Plotly JavaScript library. Full experimental acquisitions can be displayed as false color heatmaps, contour maps or 3D surfaces with a selected color palette. A filtering panel allows the application of zero-phase 2D Gaussian convolution smoothing or 2D Butterworth low pass filtering to the FSCV acquisition. For the 2D Gaussian convolution, the user selects the number of repetitions and the standard deviation of the Gaussian kernel. For the low pass filtering, the user sets the cutoff frequencies in each axis and the order of the transfer function. Current traces at selected potentials and perpendicular cyclic voltammograms can be interactively selected by the user and graphed in two distinct Cartesian axes. The extracted current traces from any of the imported FSCV files can then be calibrated *via* a calibration factor to obtain a concentration trace. The application will automatically graph the concentration trace, together with an exponential fit of the reuptake curve and calculated *t*_1/2_ of clearance. The filtered FSCV data, the calibrated curves and parametric analysis can be exported as Excel files.

The reuptake kinetic analysis is a detached application. The user can select the experimental trace used to fit the model expressed in **Equation 4**. The user can modify each of the parameters of the Michaelis Menten differential equation through the graphical interface. For each change, the application recalculates the modelled concentration profile and graphs it together with the experimental trace. Additionally, each of the Michaelis Menten kinetic parameters can be optimized using a custom-made nonlinear least-squares algorithm. As the error between the model and the experimental signal is expected to have multiple local minima, each kinetic parameter can be optimized separately providing a range of accepted values. The modelled concentration profile, as well as the optimized parameters and metrics of the goodness of fit can also be exported into an Excel spread-sheet.

### Computational Methods

Digital background subtraction was achieved by subtracting each cyclic voltammogram with an average cyclic voltammogram obtained from a user-defined section of the FSCV acquisition. Gaussian 2D convolution was carried out by separating the convolution operation into a horizontal and subsequent vertical 1D Gaussian convolution. Making use of the central limit theorem,^13, 14^ each Gaussian convolution was approximated by 3 consecutive uniform convolutions. Low pass filtering was optimized by using the conjugate symmetry property of the Fourier transform. The FSCV acquisition was mirrored at the edges before transformation to avoid low pass boundary artifacts. The 2D Butterworth filter follows the transfer function in **Equation 1**.

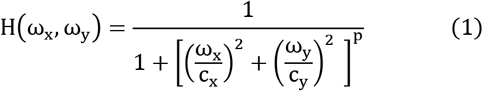

where *H*(*ω_x_,ω_y_*) is the gain at horizontal frequency *ω_x_* and vertical frequency *ω_y_, c_x_* and *c_y_* are the horizontal and vertical cutoff frequencies, and *p* is the order of the filter. The transfer function was modified from its general formulation^15^ to allow different cutoff frequencies on each axis. The bandpass frequencies have an amplification gain of 0 dB, and the cutoff values are designed as frequencies at which the gain is reduced to 50% (−3 dB).

*t*_1/2_ of clearance was estimated by fitting the exponential decay function given by **Equation 2** to the experimental evoked concentration trace after the maximum release peak.

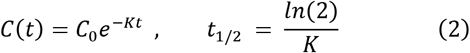

The exponential regression is initially obtained by linearizing the exponential decay function and using linear least squares. The user can manually apply nonlinear least squares to optimize the exponential fitting, in which case a custom-made stochastic gradient descent algorithm is applied to find the best fit. The stochastic algorithm uses the linear estimations as initial parameters and the root mean square error (RMSE) as cost function. The standard errors of the regression parameters were estimated from the square root of the diagonal elements of the inverted Hessian matrix. The standard error of *t*_1/2_ was estimated from the propagation of uncertainty of the K parameter,^16, 17^ as expressed in **Equation 3**.

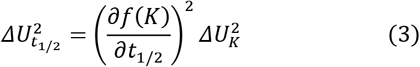

The maximum amplitude of the release of concentration profiles is detected using a custom-made peak-finding algorithm that compares each concentration value to its neighbours. The number of neighbours used is set to 20 by default and can be changed by the user. The numerical integral is calculated using the Simpson’s rule between the first point and the first intercept of the x-axis after maximum amplitude. If there is no x-axis intercept, the algorithm uses the minimum amplitude point.

The kinetics of the reuptake were estimated using one of two different mathematical models. The first one, **Equation 4**, with a single reuptake mechanism commonly used for dopamine,^18, 19^ and the second one, **Equation 5**, with two reuptake terms, previously used for serotonin.^12, 20^

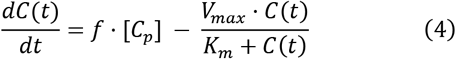

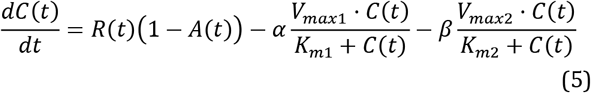

*C*(*t*) is the concentration of the neurotransmitter in the extracellular space, *f* is the frequency of stimulation, [*C_p_*] is the constant release of neurotransmitter per stimulus, *R*(*t*) is the evoked release rate of the neurotransmitter, *A*(*t*) is the fraction of occupied autoreceptors of the neurotransmitter, which works as negative feedback control, *α* and *β* are the weights of the two reuptake mechanisms, and *V_max_* and *K_m_* are Michaelis Menten reuptake parameters. The differential equations were solved by discretization of the differential term. For the second model, the initial condition corresponds to the estimated basal concentration of the neurotransmitter, provided by the user. Optimal kinetic parameters were then estimated using the previously mentioned stochastic gradient descent algorithm to fit the evoked concentration trace.

### Experimental Procedures

*The Analysis Kid* is purely a post-processing tool. In this report, we intend to describe the main functionalities of the web application. The FSCV experimental signals shown here correspond to dopamine (NAc) and serotonin (CA2 region of hippocampus) voltammetric measurements in anesthetized rodents. Experimental procedures used to acquire these signals are described extensively elsewhere.^12,21^

## RESULTS AND DISCUSSION

### Background Subtraction and Filtering of FSCV Data

FSCV is most commonly used for neurotransmitter analysis with the aim of understanding the roles and mechanisms of these chemicals in brain function and pathology. Because neurotransmission occurs on a subsecond time scale and measurable extrasynaptic neurotransmitter concentrations are low (nano to micro molar),^21, 22^ detection remains challenging. FSCV applies a continuous potential waveform to a carbon fiber microelectrode and measures the current along this potential ramp with a resolution dependent on scan rate and acquisition frequency (determined by data acquisition card). The waveform is applied at high frequency (typically 10Hz) and data are typically collected between 15-30 seconds. These data files are often presented as 2-dimensional representations of 3-dimensional data. These ‘color’ plots display the potential waveform as a function of time (y-axis) vs. data file collection time (x-axis). The current is displayed in false color. Color plots are described in more detail by Michael *et al.* ^23^ The physiological information held in these color plots is 2-fold. The potential at which redox peaks occur identifies substrates (*via* voltage vs. current (i) plots, cyclic voltammograms (CVs)), the maximum current amplitude of the peak here quantifies the analyte (current (i) vs. t).

Successful FSCV measurements necessitate several, simultaneous noise reducing/signal amplifying processes. For data acquisition, these processes include precision-fabricated electrodes, electromagnetic shielding (*via* Faraday Cage), waveform filtering and pre- and post-amplification. ^24,25^ Post-acquisition, data is digitally treated with background subtraction, filtering and multi-parametric analysis.^26^

There is no single platform that can interface with common FSCV data acquisition paradigms to simultaneously back-ground-subtract, filter and provide in-depth analysis of input data. *The Analysis Kid* aims to provide user-friendly, digital post-processing tools for FSCV data. The first step towards this is background subtraction and filtering.

Background subtraction in FSCV is necessary to remove a large, unspecific capacitive current that arises from double-layer charging on the carbon surface due to high scan rates..^27^ The idea is that over the short-term (10s of seconds) this capacitive or charging current is relatively stable. Hence, subtracting signals from a baseline will reveal the much smaller, Faradaic (electron transfer) current related to analyte concentration changes. FSCV background subtraction has been achieved in the continuous domain,^28,29^ and consists of recording the background current and subtract it from the signal at the current transducer before analog-to-digital conversion. Analog subtraction is irreversible and difficult to control as it occurs at the pre-acquisition stage. Digital, post processing methods give more visibility and control since the baseline is manually determined by a period of low activity during file collection.^30^ *The Analysis kid* enables the latter, digital background subtraction.

Filtering the digital background-subtracted signals is often critical for revealing data. Existing analysis software apply one-dimensional low pass filters to remove high-frequency noise from the cyclic voltammograms, then uniform convolutions to smooth the color plots.^31^ HDCV introduces 2-dimensional filtering by applying a one-dimensional filter to a two-dimensional mask. A two-dimensional approach removes a step, simplifying the process and rendering the data less prone to overfiltering. *The Analysis Kid* filters the high-frequency noise using two optimized approaches. Each approach is equally valid for filtering data and can be chosen depending on unique advantages and disadvantages for custom data sets.

#### 1. Two-dimensional Gaussian convolution approximated by a separable and recursive uniform convolution

Gaussian convolution better preserves the edges and overall shape of signals than uniform convolution.^32^ However the computational performance of straight Gaussian convolution is sub-optimal because the FSCV signal must be multiplied by precomputed Gaussian kernel coefficients. Specifically, the computational complexity of a two-dimensional Gaussian convolution is O(MNk^2^), with M and N being the dimensions of the color plot and k the size of the kernel. Functionally this means that because FSCV acquisitions are large data sets, filtering (depending on processing power), is not instantaneous. Our convolution algorithm maintains Gaussian smoothing properties while optimising performance by reducing the computational complexity to O(max(M, N)) - which is independent of kernel size. So while 2D FFT filtering (O(MNoln(MN))) tends to perform faster than conventional convolution operations, this is not the case for our optimized convolution algorithm, which performs approximately twice as fast as the 2D FFT filtration on the same machine and signal size.

#### 2. Two-dimensional FFT, Butterworth Low Pass Filtering

**Figure 1** illustrates the Butterworth filtering scheme. The method consists of transforming the FSCV acquisition (*in vivo* dopamine color plot, left) into the two-dimensional frequency domain (Fourier spectrum, right) (**1A**), where the y and x axes represent the vertical and horizontal frequency spectrum of the color plot. The spectrum is multiplied by the Butterworth transfer function (**1B**) using the Hadamard product (element-wise, not matrix multiplication) to scale the magnitude of the frequency components. The result of the operation is then transformed back to the temporal domain using the inverse 2D FFT algorithm (**1C**).

**Figure 1.**
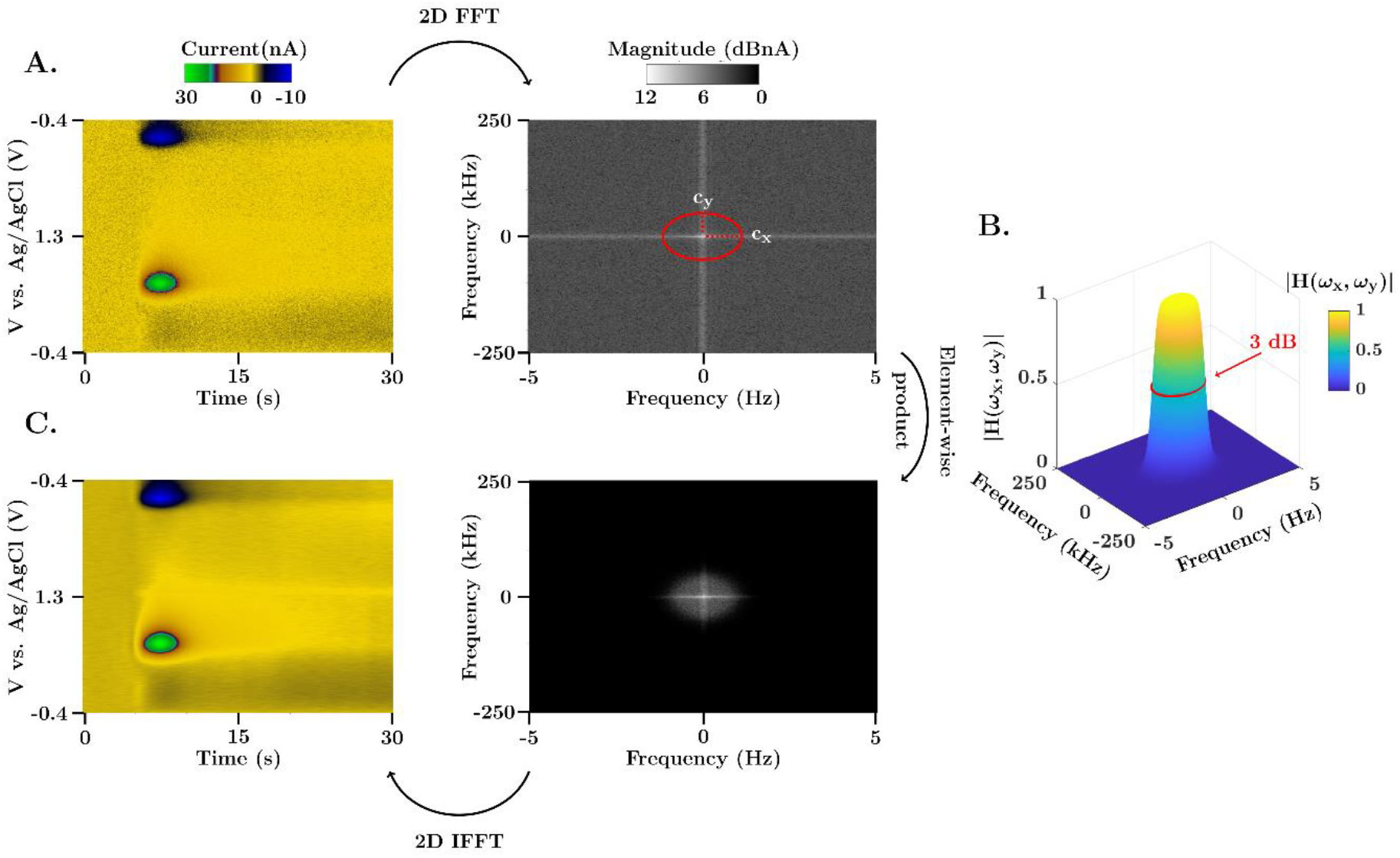
Representation of the 2D FFT low pass filtering scheme. (**A**) FSCV color plot and 2D frequency spectrum of an in vivo evoked dopamine measurements in the nucleus accumbens of a mouse brain. The range of the frequency spectrum in each axis is determined by the Nyquist rate, *f_max_* = ± *f_s_*/2. (**B**) Illustration of the 2D Butterworth filter gain (cutoff frequencies *c_x_* = 0.75 Hz and *c_y_* = 37.5 kHz, and order p = 5) for the same range of frequencies as the color plot spectrum. The 50% gain dropout (also known as 3 dB cutoff) is denoted with a red circle. The transfer function presents a uniform gain in the passband, characteristic of Butterworth filters, and a steep transition to the stopband due to the high order of the filter. (**C**) Filtered FSCV color plot and 2D frequency spectrum. The filtered spectrum is obtained as the element-wise product of the transfer function in part (**B**) and the 2D spectrum in part (**A**).

**Figure 2** provides a comparison of our two filtering methods. Here, *via* electrical stimulation (at 5 seconds) of the medial forebrain bundle in a mouse brain, dopamine is evoked. Dopamine is identified *via* the characteristic positions of redox peaks in the CV. The current signal increases immediately upon stimulation and clears within seconds afterwards. **2A** shows *in vivo* dopamine color plots (middle: raw, left: 2D convoluted, right: 2D FFT filtered). **2B** and **2C** are the i vs. time and CVs taken from the horizontal/vertical dashed crosshairs form the color plots. The unfiltered data is superimposed over the filtered data for comparison.

**Figure 2.**
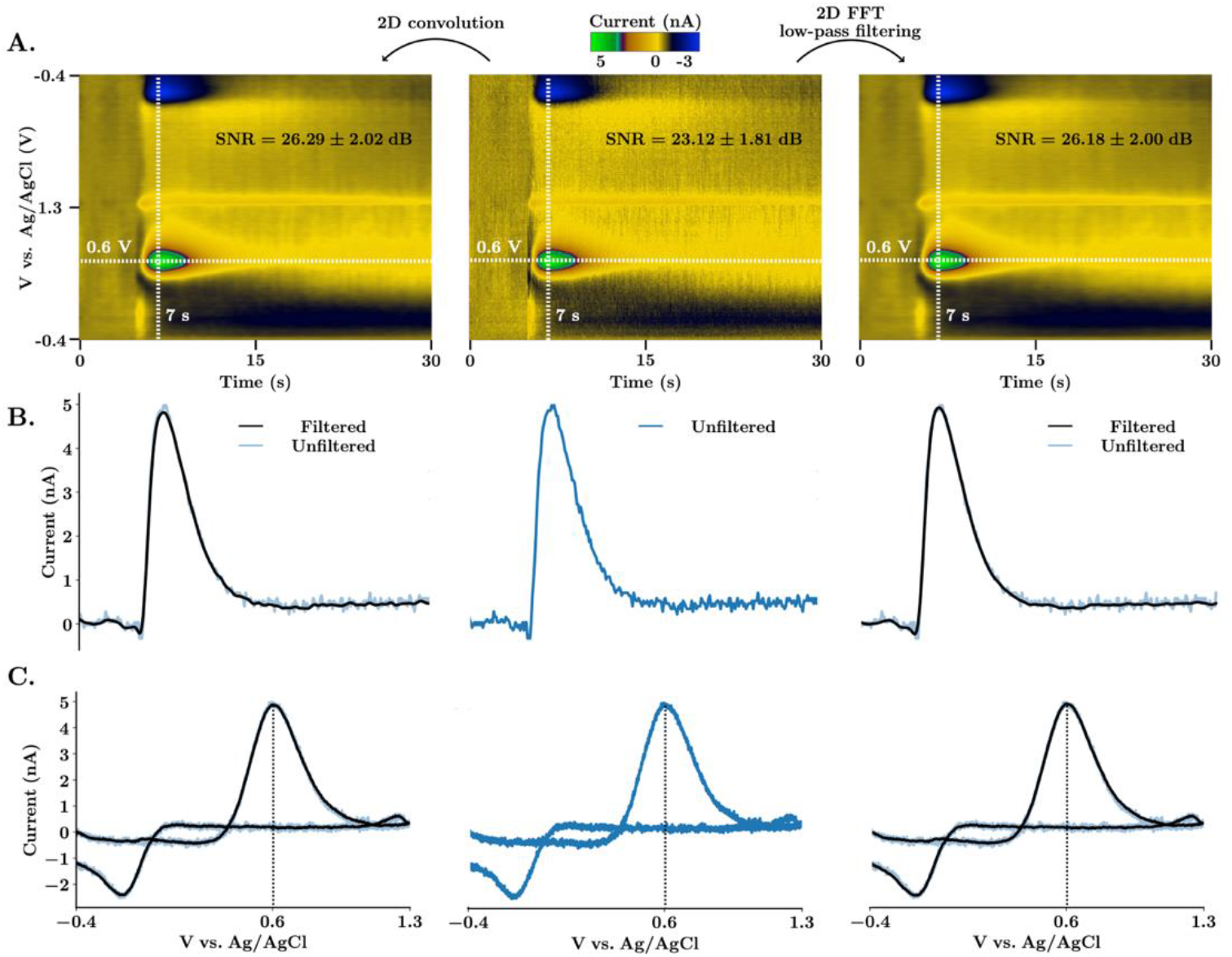
Comparison between 2D Gaussian convolution and 2D FFT low-pass spectrum filtering. (**A**) FSCV color plots of *in vivo* electrically stimulated dopamine in the nucleus accumbens of a mouse. The unfiltered color plot is shown on the center. On the left, the color plot was smoothed using 2D convolution with a Gaussian 5×5 kernel and one repetition. On the right, 2D Butterworth spectrum filtering (cutoff frequencies *c_x_* = 0.75 Hz and *c_y_* = 37.5 kHz, and order p = 5) was used instead. SNR (average ± SEM, n = 5 animals) was calculated as peak-signal-to-noise ratio (see Supporting Information). (**B**) Horizontal current traces extracted at potential 0.6 V from the respective color plot above. (**C**) Cyclic voltammograms extracted at time 7 s from the respective color plot above. Filtered signals are superimposed to the unfiltered signal to demonstrate that there are no time voltage or amplitude shifts.

Both filtering methods remove high-frequency noise and increase SNR and are zero-phase, meaning that there is no peak shift. For 2D FFT Butterworth filtering, cut-off frequencies are defined as a percentage of the maximum frequency of the signal in each direction. This allows for the selection of filter boundaries without knowledge of the frequency spectra. Each method has distinct advantages; for typical data sets, convolution is faster (*vide supra*) while filtering in the frequency domain is more intuitive and precise. Additionally, convolution in the temporal domain usually requires a trial-and-error approach. Thus, for clean, simple data convolution is more appropriate, while noisier, more complex data sets could benefit from FFT filtering. The 2D frequency spectrum representation together with the cutoff boundaries of the filter are available on the platform. As shown in **Figure 1**, the stopband of the transfer function can be designed to filter out artifactual frequency components with different cutoff frequencies for each axis. This feature is critical because the noise frequency spectra of cyclic voltammograms and current traces are different, an effect that is exacerbated because the sampling frequency used to acquire CVs is much higher than the voltage application frequency (e.g., 500 kHz vs. 10Hz).

#### Automatic Parametric Peak Analysis

Physiological information about how the concentration of an analyte changes with time is derived from color plots as i vs. time traces. The analyte concentration is directly and linearly proportional to this current,^33^ and can easily be determined *via in vitro* calibrations. A calibration factor derived from the linear portion of the calibration curve allows conversion of the signal to concentration. From these concentration signals, physiological information can be garnered from the peak amplitude and the post-stimulation clearance rate (either *via* simple *t*_1/2_ analysis of the clearance curve or a more sophisticated first-order kinetic decay described by a Michae-lis-Menten model).

*The Analysis Kid* provides a platform for simple and more complex (*i.e.* Michaelis-Menten) parametric analysis. Multiple color plot data files can be simultaneously uploaded to the platform. A single color plot is displayed, and filtering is applied (*vide supra*). *Via* an interactive selection tool, the i vs. time of interest is graphed into an embedded window and assigned a color tag. A calibration panel allows the user to apply a calibration factor to this data - the resulting concentration vs. time trace is displayed in an additional window. Here, the maximum peak concentration, *t*_1/2_ of clear-ance and area under the curve are automatically plotted. Multiple files can be treated in this way and are optionally superimposed on the i vs. t panel on the same Cartesian axes; the user can select which ones to show and hide. Each trace is assigned a tag with the name of the origin file, a color and a number. This information is shown as a tooltip when the user hovers over a current trace. The standard error of the estimate (SEE) and coefficients of the exponential fit are also provided to assess the goodness of fit. *T*_1/2_ is either obtained from a linearized exponential regression, or a nonlinear optimization of the exponential parameters.

**Figure 3** illustrates this analysis process for an *in vivo* experiment where dopamine is evoked in the same way as described above. Panel **A** is the color plot filtered with 5×5 Gaussian kernel convolution, inset into this color plot is the i vs. time extracted from the dopamine oxidation peak at 0.6 V. Panel **B and C** are the same data, calibrated for concentration. In **B**, the decay curve is fitted with a linearized exponential function (C(t)) and in **C**, the curve is fitted with a non-linear least-squares algorithm with a stopping criterion of 50 iterations per optimization. This approach better fits the data, evidenced by the lower standard error of the estimate in **C** and better R^2^ value (0.87 in **B** and 0.97 in **C**, even though the R^2^ parameter is a frail estimate of the goodness of fit of nonlinear models.^34,35^ This lower error occurs partly because a linear regression assumes the error distribution is additive in the logarithmic scale. Consequently, errors will be higher for high concentration values making a linear model less able to accurately model higher dopamine concentrations.

**Figure 3.**
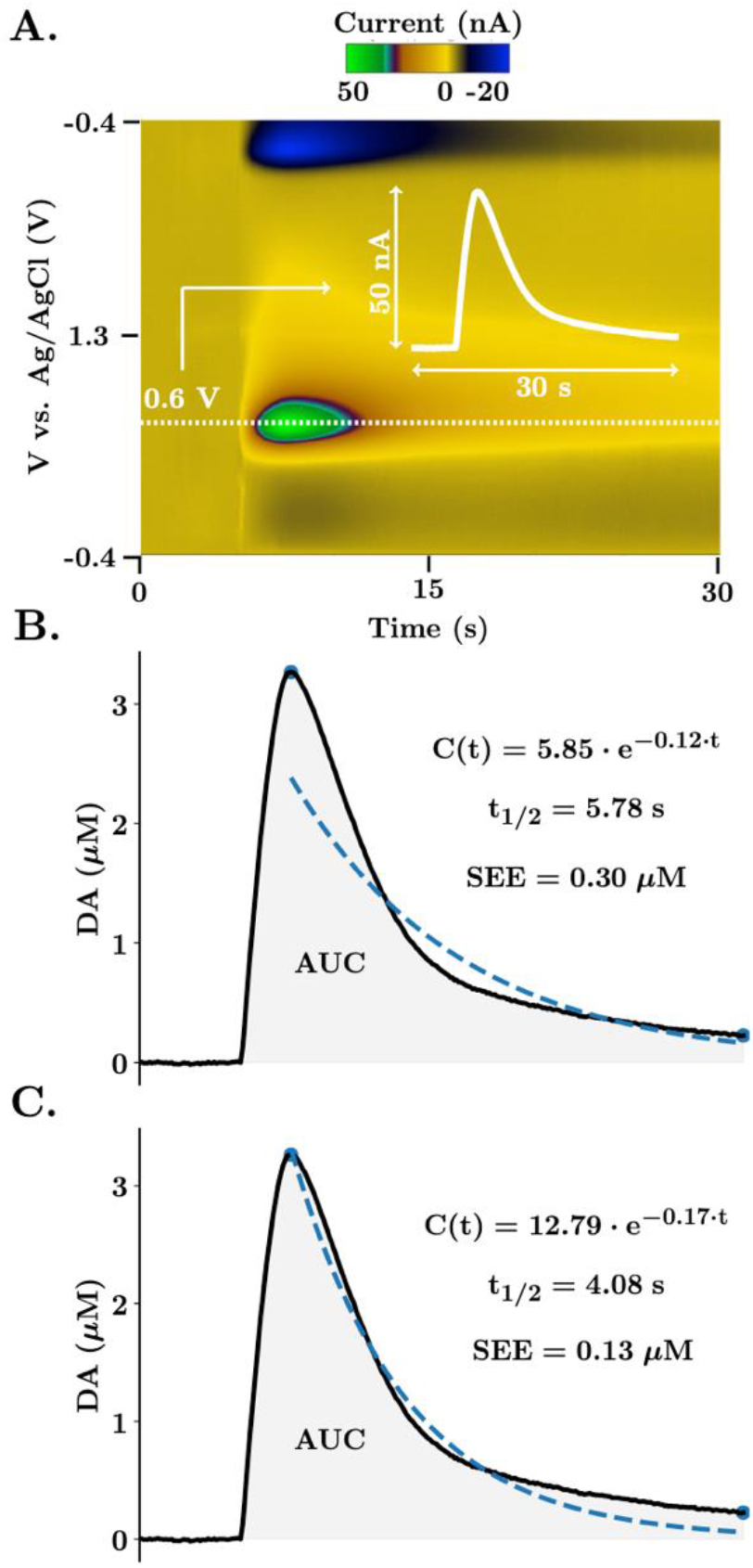
Calibration and automatic analysis in *The Analysis Kid*. (**A**) FSCV color plot of DA evoked release in the nucleus accumbens of the mouse brain. The white dotted line illustrates the extracted current trace at the faradaic potential of interest. The inset white graph shows the extracted trace. (**B, C**) Dopamine trace after calibration with a factor of 0.0625 μM/nA. Blue dots represent the maximum and minimum amplitude points detected by the algorithm. Dashed blue line represents the exponential fit, also expressed as an equation together with the half-life of the reuptake and SEE. Shaded area represents the integrated area under the curve. In part (**B**), the initial linearized exponential regression is provided. In part (**C**), the nonlinear least-squares algorithm is applied for 50 iterations.

#### Analysis of Reuptake Kinetics

Michalis-Menten models are employed to analyze reuptake curves because transporter/substrate interactions have first-order reaction kinetics. The M-M equation, **Equation 6** below, describes the velocity of reaction (v) as a function of substrate concentration and two constants that define the system. *V_max_* is the maximum rate of reaction, the higher *V_max_* the more efficient the transporter. *K_m_* is the substrate concentrate at 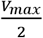 and measures the capacity of the system. The lower the *K_m_*, the higher the affinity of the transporter for the substrate.

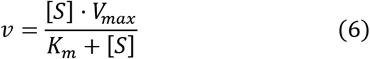

Thus, modelling reuptake curves with M-M informs about the efficiency and affinity of transport systems to be established. Different M-M models have been developed to fit FSCV concentration traces, including general models incorporating one^19,36,37^ or two^12,20^ reuptake mechanisms, and circuit-level models designed for multiple neurotransmitters.^38^ All of these models are differential equations. Most commonly computational software (such as MATLAB) is utilized to fit the solution of the differential equation to the experimental data. This approach, while effective, requires a high level of computational and mathematical expertise. In *The Analysis Kid*, we created a straightforward estimation of reuptake kinetics from two general models with one and two reuptake mechanisms (**Equation 4** and **Equation 5**, used previously to model dopamine and serotonin traces, respectively).

**Equation 4** is a simple model that has been extensively used to model dopamine curves with only 3 variables (*V_max_*, *K_m_* and [*C_p_*]), describing the rate of change of analyte (dopamine) as a function of dopamine released per stimulation pulse minus a single M-M term.^19^ However, this equation was not successful in modelling serotonin curves because of two additional features of the experimental traces. These additional experimental features arise from a prolonged autoreceptor effect (1-A(t)) that offsets release, and a second M-M term to reflect Uptake 1 and 2 mechanisms. These multiple reuptake mechanisms have been described for some time for serotonin. The notion is that at low serotonergic activity, the serotonin transporters, that are localized to the synaptic region, uptake serotonin with high affinity but low capacity, this mechanism is called Uptake 1. At times of increased activity (above a certain concentration threshold), other monoamine transporters such as norepinephrine and organic cation transporters team up to uptake serotonin. This Uptake 2 mechanism clears serotonin with low affinity (because the transporters are not specific to serotonin) and high capacity (but there are several different types). α and β coefficients allow different weights to be assigned to each reuptake component dependent on local tissue architecture and a concentration threshold defines at which point Uptake 2 is activated.^20^ The dopamine model assumes a basal level of 0, while the serotonin model assumes a basal level of 20-80 nM based on previous work. First, we found that serotonin falls below baseline by up to 10s of nM after stimulation - effect that we attributed to autoreceptors,^20^ and second from actual basal level measurements of serotonin with a new technique FSCAV.^39^

The analysis is partially automatic. For dopamine, input of stimulation frequency and the number of pulses is sufficient for an initial estimation of the modelled trace. Then, the user can apply our optimization algorithm - a stochastic gradient descent algorithm that iterates over user-fixed intervals of *K_m_, V_max_* and [*C_p_*] to improve the model fitting. For serotonin, the release and autoreceptor term are modelled interactively in the user interface and are not defined by constants. The release term can take any shape, which is necessary to fit the experimental data arising from the complex release and auto-inhibition interaction. Standard values of *K_m_* and *V_max_* for Uptake 1 and 2 are the starting points for modelling the two reuptake curves. Basal levels, α and β coefficients and a threshold concentration for Uptake 2 (β) are user inputted and the fitting is as for dopamine above. When multiple parameters are optimized, the algorithm will iterate them in succession. The number of iterations and learning rate of the optimization process can also be modified by the user.

**Figure 4** shows how stereotypical *in vivo* dopamine (**A**) and serotonin (**B**) traces are modeled with *The Analysis Kid.* **Ai** and **Bi** contain the color plots, filtered with a 5×5 kernel con-volution. **Aii** and **Bii** show the model fits, and **Aiii** and **Biii** show how the release (and autoreceptor for serotonin) terms change with time. For dopamine, the M-M parameters of the single reuptake (*V_max_* = 1.45 *μM/s,K_m_* = 0.39 *μM*, [*C_p_*] = 0.04 *μM*) agree with those from previous studies in the NAc of rats.^40^ The release term (**4Aiii**) is modelled as a constant release of neurotransmitter during stimulation with only one variable parameter([*C_p_*]), and the model fits the experimental data with a low deviation from the experimental trace. For serotonin, an experimental signal with two different reuptake curves was analyzed. The slow reuptake mechanism (*V_max_* = 14.59 *nM/s,K*_*m*1_ = 5.09 *nM*) always takes effect (*α* = 1), while the fast reuptake mechanism (*V*_*max*2_ = 642.38 *nM/s,K*_*m*2_ = 221.63 *nM*) only takes effects when the experimental trace reaches a concentration threshold (*β* = 0.03, *threshold* = 22 *nM*). The release and autoreceptor terms are manually modelled to fit the shape of the experimental signal. The optimised kinetic parameters and overall shape of the release and autoreceptor terms agree with previous work.^20^ Thus, our application provides an analogous yet straightforward method to model the reuptake kinetics.

**Figure 4:**
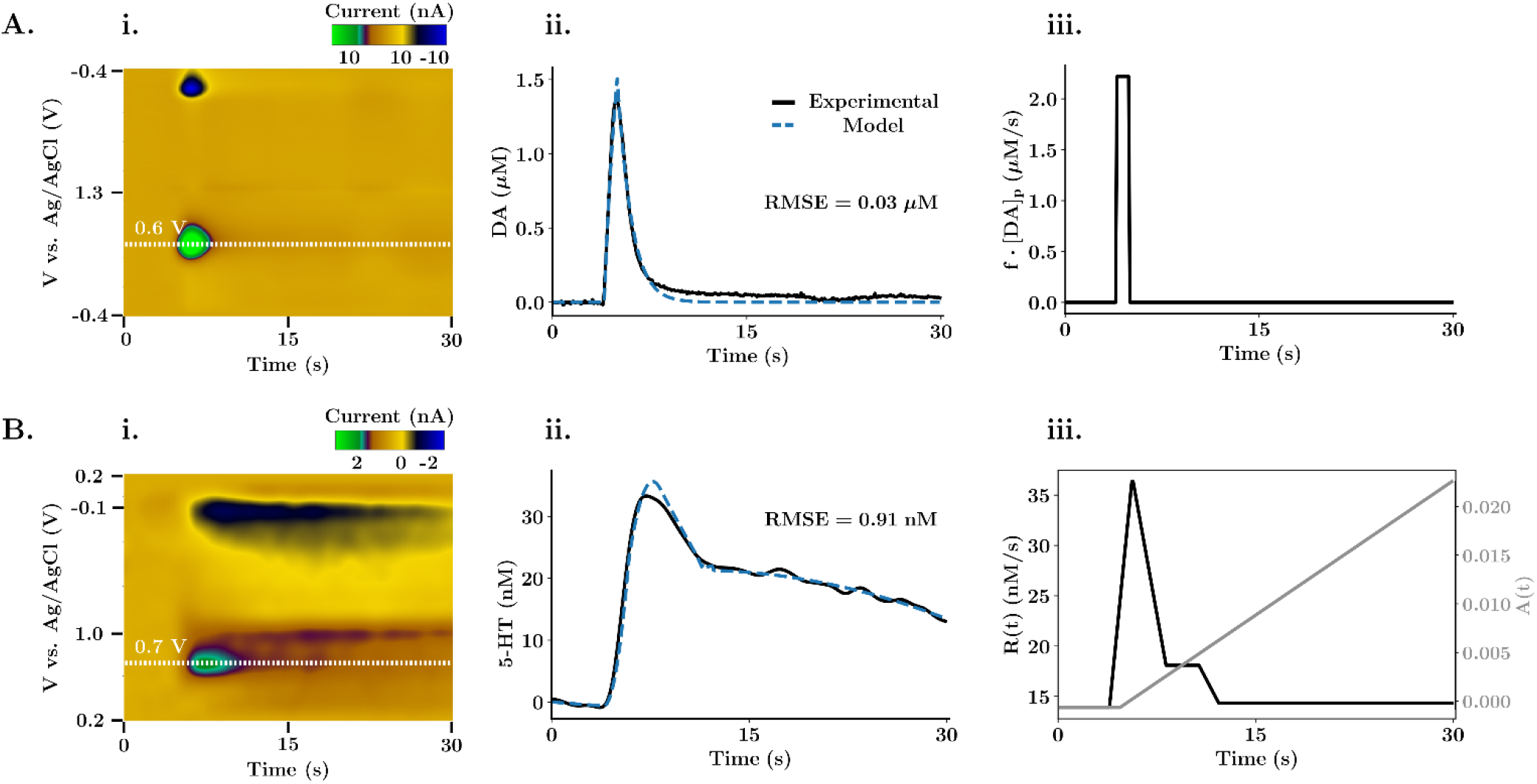
Michaelis-Menten reuptake kinetics fitting. (**A**) Modelled one reuptake kinetic analysis for DA evoked release in the NAc. (**B**) Modelled dual reuptake kinetic analysis for 5-HT evoked release in the CA2 region of a mouse hypothalamus. (**Ai, Bi**) FSCV color plot. The white dotted line shows the extracted current trace, calibrated with a factor of 0.0625 μM/nA for dopamine and 20.20 nM/nA for serotonin. (**Aii, Bii**) Experimental release trace and corresponding modelled trace obtained with the kinetic analysis. The root mean squared error was used by the optimization algorithm to assess the goodness of fit. Each of the optimization used 2000 iterations at a relative learning rate of 0.001 for each of the variables in the equation, (**Aiii, Biii**), Release term (and autoreceptor term for **Biii**) of the modelled differential equation.

## CONCLUSIONS

For filtering and multi-modal analysis of FSCV signals several custom platforms are available. However, no easily accessible software has the capacity for all these features. In this work, we introduced *The Analysis Kid*: a free, open-source web application. We showed that multi-platform file formats could be imported for visualization of FSCV color plots with background subtraction. We discussed key features that enabled calibration and parametric analysis (maximum amplitude, AUC and *t*_1/2_) of the signal. Finally, we highlighted a key feature of *The Analysis Kid,* namely semi-automatic fitting of data with Michaelis Menten kinetic models for *in vivo* dopamine and serotonin traces. *The Analysis Kid* aims to simplify and broaden FSCV analysis significantly.

## ASSOCIATED CONTENT

### Supporting Information

This material is available free of charge via the Internet at http://pubs.acs.org. This includes a detailed video tutorial of the web application.

## AUTHOR INFORMATION

### Author Contributions

SM developed the software, SD contributed to the development of the reuptake kinetics application, CW and SB collected *in vivo* data, PH and SM wrote the manuscript. The manuscript was written through contributions of all authors. / All authors have given approval to the final version of the manuscript. / ‡These authors contributed equally.

### Notes

The authors declare not competing financial interest.

## ACKNOWLEDGMENTS

The authors wish to thank members of the Hashemi Lab and Pavithra Pathirathna for helpful discussions and feedback. Additionally, the authors are grateful to Julie Hoang for capturing and editing the instructional video.

## ABBREVIATIONS

FSCV: fast-scan cyclic voltammetry
SNR: signal-to-noise ratio
5-HT: 5-hydroxytryptamine
DA: dopamine
FFT: fast Fourier transform
RMSE: root mean square error
SEE: standard error of the estimate
SEM: standard error of the mean

